# scDeepHash: An automatic cell type annotation and cell retrieval method for large-scale scRNA-seq datasets using neural network-based hashing

**DOI:** 10.1101/2021.11.08.467820

**Authors:** Shihao Ma, Yanyi Zhang, Bohao Wang, Zian Hu, Jingwei Zhang, Bo Wang

## Abstract

Single-cell RNA-sequencing technologies measure transcriptomic expressions, which quantifies cell-to-cell heterogeneity at an unprecedented resolution. As these technologies become more readily available, the number of scRNA-seq datasets increases drastically. Prior works have demonstrated that bias-free, holistic single-cell profiling infrastructures are essential to the emerging automatic cell-type annotation methods. We propose scDeepHash, a scalable scRNA-seq analytic tool that employs content-based deep hashing to index single-cell gene expressions. scDeepHash allows for fast and accurate automated cell-type annotation and similar-cell retrieval. We also demonstrate the performance of scDeepHash by benchmarking it against current state-of-the-art methods across multiple public scRNA-seq datasets.

## 1 Introduction

Single-cell RNA sequencing technology quantifies transcriptomic expressions at single-cell level, leading to discoveries rooted in cell-to-cell heterogeneity. Cell-type annotation, or assignment of cell type labels to single cells, is an important step in analysing the RNA sequencing data (Lähnemann et al. [2020]). With its wide and mature applications, scRNA-seq datasets have become increasingly and readily available in a larger scale. The scale of scRNA-seq datasets has increased significantly in sample sizes, dimensions of features (genes), the number of heterogeneous cell populations and inter-correlation between cell populations. Such explosive emergence of large-scale scRNA-seq data makes manual annotation no longer desirable due to the need of labor-intensive efforts, and demands for a more reliable and efficient automated cell-type annotation method for biological downstream analysis.

Existing cell-type annotation methods can be classified into three general categories (note that these categories are not mutually exclusive and can overlap for some methods): annotation based on known marker genes following clustering; annotation based on similarity between the query dataset and the labeled reference dataset; supervised learning that labels query cells using a classifier trained from a labeled reference dataset. Yet, these methods cannot be easily adapted to annotate large-scale scRNA-seq datasets. Marker gene-based methods are labor intensive in practice and the results can have high variation; similarity-based methods usually suffer from the trade-off between accuracy and speed; machine learningbased methods can have inconsistent performance on different datasets. We discuss the details of existing methods in Section 2.

Through compression of high-dimensional data into much lower one, hashing has proven its high scalability for the ultra-fast data look-up speed and efficient data storage in large-scale data scenarios. Since the core of hashing algorithm - the hash function - is highly data-dependent, the recent emerging ‘learning-to-hash’ methodology aims at supervisingly learning it so that the nearest neighbor search result in the hash code space is as close as the search result in the original space (Wang et al. [2018]). In this work, we propose scDeepHash - a ‘learning-to-hash’ method that can encode distinguishable single-cell gene expressions into short, binary hash codes and map them to their corresponding ‘hash buckets’, each representing a distinct cell type as an cell anchor. With well-trained hash function to encode high-dimensional scRNA-seq data into binary code, scDeepHash can handle large-scale scRNA-seq datasets by accurately, efficiently annotating and retrieving single-cell data.

Our main contributions are summarized as follows: **(1)** our high-quality hash codes well preserve the similarity information within the same cell type in the original gene expression space; **(2)** the short hash codes enable efficient binary computation for cell similarity; **(3)** in addition to cell-type annotation, scDeepHash can also well perform similar-cell retrieval tasks. That is, given a query cell, scDeepHash can reliably and efficiently search and retrieve the most similar cells within a database. To our best knowledge, this is the first time the idea of ‘learning-to-hash’ is applied to cell-type annotation and similar-cell retrieval. We demonstrate that scDeepHash achieves comparable performance or outperforms other state of the art on accuracy and speed for both cell-type annotation and similar-cell retrieval on extensive large-scale scRNA-seq datasets.

## 2 Related Works

Existing methods for cell-type annotation can be categorized into three general but not mutually-exclusive types: marker gene-based methods, similarity-based methods and machine learning-based methods.

**Marker gene-based methods** utilize biological prior knowledge for interpretable annotations. scRNA-seq data, after dimension reduction and clustering (Butler et al. [2018], Luecken and Theis [2019]), are manually annotated on cluster level by referencing the highly expressed genes with public cell-type marker gene databases such as CellMarker and PanglaoDB (Zhang et al. [2018], Franzén et al. [2019]). An alternative approach is to incorporate the prior knowledge with a statistical inference framework. For example, SCINA (Zhang et al. [2019a]) operates at cluster level by fitting a bimodal distribution to the expression of marker genes. However, the clustering-based gene selection is very sensitive to the clustering methods, choice of hyperparameters, and the number of cells in datasets, which can cause large variation in the downstream analysis (Alquicira-Hernandez et al. [2019], Kiselev et al. [2019]). Another example is CellAssign (Zhang et al. [2019b]) that includes a marker gene-based matrix within a probabilistic graphical model to perform annotation at single-cell level, which avoids the clustering step. However, its annotation performance is still largely dependent on the choice of canonical cell markers, which is often complex and labor intensive in practice (Alquicira-Hernandez et al. [2019], Kiselev et al. [2019]).

**Similarity-based methods** assign cell labels by measuring the similarities between query cells and reference datasets, which can be categorized into two types: cluster-to-reference, or single cell-to-reference. Examples like CIPR (Ekiz et al. [2020]) and ClustifyR (Fu et al. [2019]) fall into the former category, which unavoidably suffers from the limitations of clustering; for the latter category, notable tools include scmap (Kiselev et al. [2018]), SingleR (Aran et al. [2019]), and scMatch (Hou et al. [2019]), which generally leverage statistical metrics (like Spearman and Pearson correlation) to determine the similarity between each query cell and reference public datasets like Blueprint (Stunnenberg and Hirst [2016]) and FANTOM (Lizio et al. [2016]) for annotation. However, such methods can hardly balance the trade-off between accuracy and speed. Exceptionally fast inference can be achieved when each query cell is compared with the approximate referenced gene expression (like referenced cluster centroid in scmap-cluster (Kiselev et al. [2018])), but annotation accuracy will be significantly impaired; comparing cell to cell and then iteratively performing scoring aggregation per label like SingleR (Aran et al. [2019]) can improve the annotation accuracy, but it is at the big cost of annotation speed.

**Machine learning-based methods** aim at overcoming the intrinsic variation of data and noise to find the invariant latent features through learning the model distribution of labeled datasets. Methods built on top of traditional classification methods such as Random Forest (Cahan et al. [2014], Tan and Cahan [2019]), KNN (Wang et al. [2019], Lin et al. [2020]) and SVM (Wagner and Yanai [2018], Alquicira-Hernandez et al. [2019]) can achieve decent results, but they are limited to application to cell-type annotation only. Recent works developed more complex deep learning architectures to increase models’ capability. For example, scVI (Lopez et al. [2018]) uses a hierarchical Bayesian model based on a modification of Variational Autoencoder (VAE) to learn the invariant latent features, but its annotation performance will encounter sharp decrease on deeply annotated or highly inter-correlated datasets (Abdelaal et al. [2019]); MARS (Brbić et al. [2020]) applies meta learning through joint annotated and unannotated experiments to learn the shared feature among similar cells. However, users cannot perform annotation until the successful joint training, which sharply slows down its annotation speed.

## 3 Methods

Suppose we are given the high dimensional single-cell RNA-seq data as 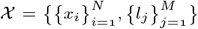, where 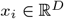 denotes each cell data in total of *N* cells, and *l_j_* ∈ *L* denotes the corresponding cell label in total of *M* different cell types. Based on *L*, we generate *K*-bit cell anchors on Hamming space: 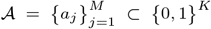, each representing an heterogeneous cell type. The objective of scDeepHash is to learn a nonlinear hash function that can map the input RNA-seq data to their corresponding cell anchors, where each cell is encoded as a *K*-bit hash code *h_i_* ∈ {0, 1}^*K*^. As shown in Figure 1, scDeepHash consists of three sequential phases: (1) cell anchors generation, (2) hash function training, and (3) cell annotation or retrieval.

**Fig. 1.**
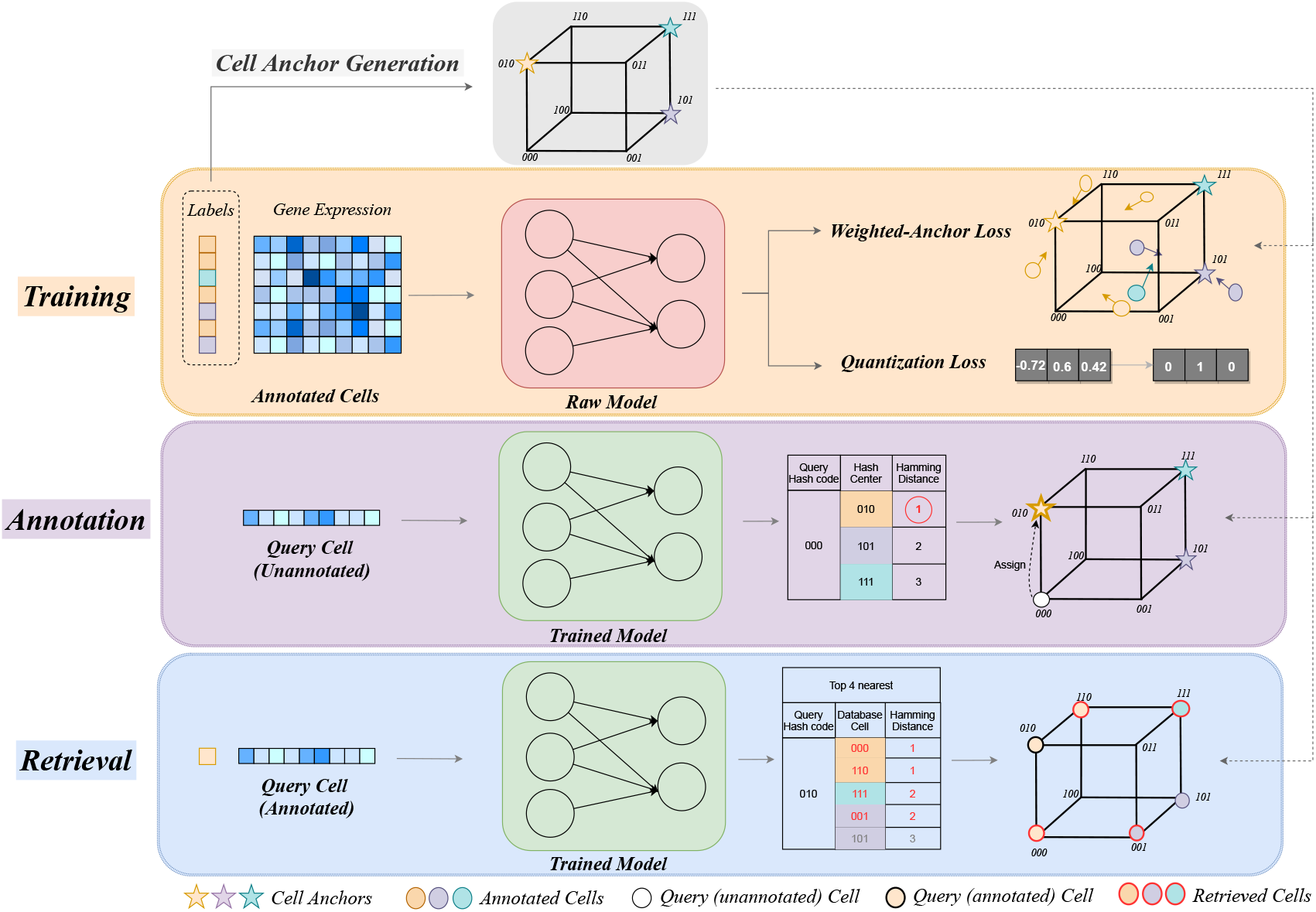
The overview of scDeepHash. scDeepHash consists of three sequential phases: **(1)** cell anchor generation (in grey): from the annotated cells’ labels, cell anchors are generated leveraging properties of Hadamard Matrix (Sec. 3.1). The generated anchors will be used as basis for the training phase and annotation phase. **(2)** Hash function training (in orange): scRNA-seq dataset is used to supervisingly train the deep learning model for optimizing the targeted hash function constrained with two specially-designed losses: weighted cell-anchor loss and quantization loss (Sec. 3.2). **(3)** Cell annotation (in purple) or retrieval (in blue): for annotation, unannotated scRNA-seq data will be fed through the trained model to generate the query cell’s hash codes, which are used for cell-type annotation based on closest-hash center strategy (Sec. 3.3); for retrieval, annotated scRNA-seq data will also be passed through the trained model to generate the hash codes, which are utilized for retrieval of the top-*k* similar cells in database (Sec. 3.3).

### 3.1 Cell Anchors Generation

Inspired by hashing methodology, we propose to introduce cell anchors (like hash buckets in the hashing community) to represent heterogeneous cell types in the Hamming space. To better distribute cell data of different types to cell anchors in the Hamming space, cell anchors should be mutually distant to preserve the dissimilarity of different cell types. The distance between two cell anchors *a_i_*, *a_j_* can be defined using Hamming distance *D_H_* (*a_i_*, *a_j_*), which denotes the number of different bits between two hash codes.

Thus, we leverage the properties of Hadamard matrix for our cell anchor generation. For a *K* × *K* Hadamard matrix 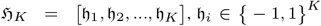, the pairwise Hamming distance of any two column vectors is exactly 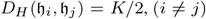, which is desirable for the purpose of cell anchors generation. Thus, cell anchors can be obtained by sampling the column vectors and then converting them into Hamming space {0, 1}^*K*^ by simply replacing all the −1 with 0. We use *Sylvester’s construction* (Yarlagadda and Hershey [1997]) to construct a *K*-dimensional Hadamard matrix:

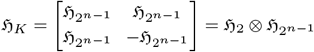

where 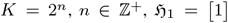, and ⊗ denotes the *Kronecker product* (Loan [2000]). We choose the n such that 2^*n*-1^ < *M* < 2^*n*^ to guarantee that we have enough column vectors 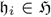 in the Hadamard matrix from which we directly sample the cell anchors 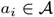.

Aside from pre-defined cell anchors, we have also tried to jointly learn the cell anchors from high-dimensional raw data with the hash function. As further discussed in Section 4.3, predefining cell anchor is a better choice due to its better performance and simplicity. Thus, the first step of scDeepHash is to generate pre-defined cell anchors.

### 3.2 Hash Function Training

Using deep learning to learn hash functions, also known as deep hashing, is a well-established methodology in the hashing community. As shown in DHN (Zhu et al. [2016]), DCH (Cao et al. [2018]) and HashNet (Cao et al. [2017]), the loss function is usually formulated as Maximum a Posterior (mAP) estimation, which is then split into two components using the Bayes’ theorem. The first component is the likelihood function, which, in our context, aims at maximizing the probability of finding the corresponding cell anchor given a cell hash code. The second component is the prior distribution over hash codes, which aims at enforcing minimum information loss when quantizing continuous codes into discrete binary hash codes to converge to corresponding cell anchors. In scDeepHash, we formulate the two losses as *Weighted Cell-Anchor Loss* and *Quantization Loss*.

#### Weighted Cell-Anchor Loss

*L_CA_*. Define the generated hash code set as 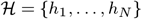, and the cell anchor set 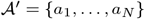, where *h_i_* and *a_i_* are the learned hash code and the corresponding cell anchor for cell *i*, respectively (We use 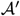 here for better illustration of the equations). The purpose of *Weighted Cell-Anchor Loss* is to aggregate the generated hash codes of cell data around their corresponding cell anchors so that the difference between them is minimized. Since all the cell anchors are binary vectors of 0 and 1, we adopt the Binary Cross Entropy (BCE) loss to penalize the differences bit by bit. The presence of rare cell types may induce an imbalanced representation of the available cell samples, which leads to poor annotation performance for the rare cell types. Thus, we introduce a weight coefficient based on the effective number of cells in each cell types to handle imbalanced data training (Cui et al. [2019]):

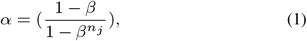

where *β* is a hyper-parameter ∈ [0, 1), and *n_j_* is the number of cell samples of j-th cell type. We then calculate the weight coefficient for each cell *i* using (1) and obtain *α* = {*α*_1_, …, *α_N_*}. Assuming the hash codes and the cell anchors are k-bits, the *Weighted Cell-Anchor Loss* is:

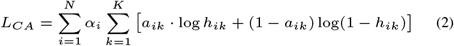

#### Quantization Loss

*L_Q_*. Since discrete optimization of the generated hash codes is challenging, applying continuous relaxation to the binary constraints during training is a standard practice among deep hashing methods. Quantization loss is needed to minimize the error caused by continuous relaxation to make sure the model can learn high-quality hash codes. To put it in biological context, we want minimum loss of cell similarity information when we turn continuous hash codes into binary hash codes. Inspired by DCH (Cao et al. [2018]), we used a Cauchy distribution prior to control the quantization error, which is defined as:

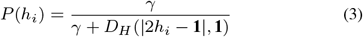

where *γ* is the scale parameter of Cauchy distribution and **1** is a vector of 1’s. The benefit of Cauchy distribution is that it can penalize significantly when similar pairs having large Hamming distance, which is exactly what we want in 3 to make the learned hash codes as close as possible to discrete {0, 1}^*K*^. Formally, the quantization loss can then be defined as:

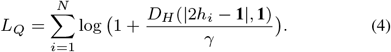

Finally, combining (2) and (4), the final target loss function of scDeepHash is

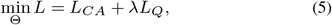

where Θ is the parameter set for *L*, and λ is a hyper-parameter to calibrate the weight of quantization.

### 3.3 Cell Annotation & Retrieval

With a well-trained model that can extract high-dimensional cell features (genes) into distinguishable binary hash code, scDeepHash can thus perform both cell-type annotation and similar-cell retrieval in an accurate and efficient way.

#### Annotation

Each query cell is passed through the network to generate its corresponding hash code. After comparing the pairwise Hamming distance between the cell’s hash code and each cell anchor, we choose the closest cell anchor as the query cell’s assignment (Figure 1), which we call the Closest Cell Anchor **(CCA)** strategy. It is simple but very efficient since its computation complexity is **O(M)**. This can ensure that the annotation can be applied to increasingly large-scale unannotated datasets without sacrificing the annotation speed.

#### Retrieval

All the cell hash codes within the database are first split into different ‘hash buckets’ based on their corresponding cell anchors. During retrieval, the generated hash code of each query cell is used to compute the closest cell anchor, similar to the cell annotation process. Then, within the selected cell anchor, *k* nearest-neighbor strategy (Figure 1) is used to fetch the top k-nearest cells, where *k* is of users’ choice. If *k* is larger than the size of the selected bucket, we move to the next closest cell anchor and repeat the same process. In general, *k* ≪ *N/M*, and the retrieval computation complexity is **O(M + N/M)**, where N/M is the average number of cells in all cell anchors.

### 3.4 Hyper-parameters Settings

Our neural network has five fully connected layers. Rectified linear unit (ReLU) function is used as activation function for all the layers, with an extra sign function to quantize the final network’s outputs. The network is trained using stochastic gradient descent (Adam optimizer) with learning rate of 1 × 10^−5^, weight decay of 5 × 10^−4^ and batch size of 128. The regularization constant λ is settled to 1 × 10^−4^ while the *β* in weight coefficient is set to 0.9999 through grid search. Number of hashing bits *K* is determined based on the number of cell types M, as mentioned in Section 3.1. The entire code base is implemented using PyTorch Lightning and is available at *https://github.com/bowang-lab/scDeepHash*.

## 4 Results

To demonstrate the superiority of scDeepHash in both cell annotation and retrieval, we have selected a variety of scRNA-seq datasets listed in Table 1. These datasets can well represent different cell data ‘complexity’: (1) the number of cells, (2) the number of features (genes), (3) the annotation levels, (4) the correlation between cell populations, and (5) different protocols, each representing a different level of challenge to learn the underlying hash function mapping. We perform the same pre-processing steps as (Abdelaal et al. [2019]).

**Table 1.** The configurations of datasets used for cell-type annotation and similar-cell retrieval experiments.

| Dataset | $n_{\text{cells}}$ | $n_{\text{genes}}$ | $n_{\text{cell types}} (>10 \text{ cells})$ | Description | Protocol | Reference |
| --- | --- | --- | --- | --- | --- | --- |
| Xin | 1,449 | 33,889 | 4(4) | Human pancreas | SMARTer | Xin et al. [2016] |
| Baron Human | 8,569 | 17,499 | 14(13) | Human pancreas | inDrop | Baron et al. [2016] |
| Muraroa | 2,122 | 18,915 | 9(8) | Human pancreas | CEL-Seq2 | Muraro et al. [2016] |
| Segerstolpea | 2,133 | 22,757 | 13(9) | Human pancreas | SMART-Seq2 | Segerstolpe et al. [2016] |
| TM | 54,865 | 19,791 | 55(55) | Whole Mus musculus | SMART-Seq2 | Nicholas et al. [2018] |
| AMB92 | 12,832 | 42,625 | 110(92) | Primary mouse visual cortex | SMART-Seq v4 | Tasic et al. [2018] |
| Zheng 68K | 68,579 | 20,387 | 11(11) | PBMC | 10X CHROMIUM | Zheng et al. [2017] |

### 4.1 Performance on Cell-Type Annotation

We select top six state-of-the-art methods for benchmarking according to the annotation performances on both accuracy and speed (Abdelaal et al. [2019]), including SingleCellNet (Tan and Cahan [2019]), SingleR (Aran et al. [2019]), ACTINN (Ma and Pellegrini [2019]), CaSTLe (Lieberman et al. [2018]), scVI (Lopez et al. [2018]) and SVM.

#### Intra-dataset annotation accuracy

We use the following 5 scRNA-seq datasets with different complexities for benchmarking intra-dataset annotation: Xin, BaronHuman, TM, AMB92 and Zheng 68K. Xin and BaronHuman represent typical-sized scRNA-seq datasets (around 1.5k to 9k cells), while TM and Zheng 68K represents large scRNA-seq datasets (50k+ cells); both Xin and AMB92 have relatively high feature dimensions (33k+ genes), and AMB92 also represents a much more granularly annotated dataset with 92 post-processed cell populations; Zheng 68K is the most ‘challenging’ dataset for hash function learning because the samples are highly inter-correlated (Abdelaal et al. [2019]). All models are trained using 5-fold cross-validation on each dataset separately. For each dataset, we measure the annotation accuracy based on median F1-score (the median of F1-scores across all cell types).

Figure 2(a) shows that the annotation accuracy of all methods decreases on datasets with increasing number of cells and features, annotation levels, and correlation. On relatively ‘easy’ datasets like Xin and Baron Human, all methods perform relatively well. However, when applying on TM, competing methods start to experience a notable drop, with SingleR exhibiting a sharp drop below 90%, and scVI even to 0% (beyond model’s capacity, and thus not display). In this case, scDeepHash’s accuracy is only second to SVM with a negligible gap (97.3% compared with 98.0%). This shows that when scaling to cell datasets with much larger sample size and gene dimension, scDeepHash can still maintain a good performance on cell-type annotation. This resonates with the contribution of Weighted Cell-Anchor loss and Quantization loss: high-dimensional and continuous features can be ‘compressed’ to the ones in discrete hash code while still maintaining the most distinguishable characteristics. As a learning-based method, more cell samples can better help learning the hashing mapping.

**Fig. 2.**
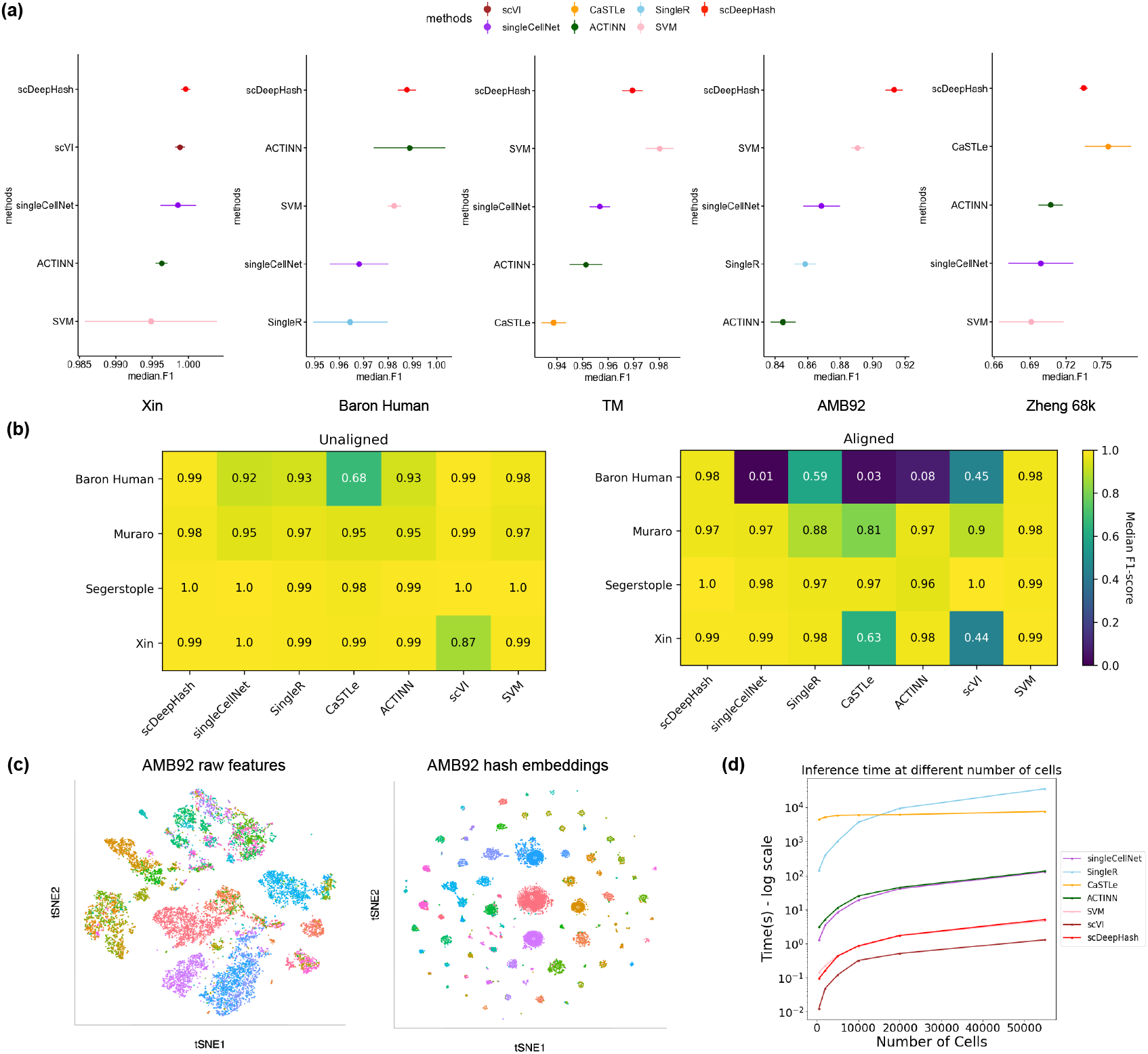
Annotation experiments. **(a)**: The intra-dataset annotation performance measured by Median F1-score on the five datasets of all benchmarked methods in boxplot. Each method is tested using the 5-fold cross-validation on each dataset. Aside from scDeepHash, for each dataset we only visualize the top-4 best performing methods. **(b)**: The inter-dataset annotation performance measured by Median F1-score across pancreatic datasets in heatmap. The row labels indicate which dataset was used as the test set while the other three datasets were used as training set. **(c)**: The prior- and post-hashing cell data in t-sne visualization of scDeepHash on AMB92. **(d)**: Measured in seconds, the annotation speed on simulated TM dataset across all selected methods.

On the AMB92 dataset with 92 cell populations (after filtering) and 42k genes, scDeepHash ranks the best by a relatively big margin, surpassing the second best SVM for around 3% (92.0% to 89.1%). This shows that comparing with other state-of-the-art methods, scDeepHash performs well on handling high-dimensional cell datasets with abundant cell populations. The t-sne visualization of the raw and hashed cell data distribution in Figure 2(c) further verifies such a superior performance in an intuitive way: each cell anchor represents a different cell type individually, aggregates its corresponding hashed cell data into separate, isolated clusters on the hamming space. Based on the underlying methodology, the superiority of scDeepHash’s annotation accuracy can be more evident with increasing number of cell populations and gene features.

On the Zheng 68k dataset, all methods encounter a sharp decrease in annotation accuracy. After analyzing the annotation results, we find that the decrease in the scDeepHash annotation accuracy is mainly due to the wrong annotation of the CD4+ T population, of which all the 4 sub-populations have below 40% of correct annotation across all methods. This resonates with the high inter-correlation among the Zheng 68k’s cell populations, which makes it very difficult to annotate. Here, scDeepHash ranks second to CaSTLe with a gap of ~1.6%. This is an interesting finding because the performance of CaSTLe is much worse on the other four benchmarked datasets. CaSTLe performs very heavy data pre-processing steps to extract the final 100 features for training on XGBoost, and one of the steps is to remove the highly inter-correlated features (Lieberman et al. [2018]). We suspect that such step is especially helpful in datasets with abundant highly correlated genes, but for those datasets with less inter-correlation, simply extracting 100 genes of high-dimensional cell data (~20k original genes) may lose too many distinguishable features and lead to poor annotation performance.

It is worth to note that scDeepHash annotation accuracy is robustly stable on the five selected datasets, with the biggest fluctuation below 1.2%, while the other methods exhibit high variance on different folds within a single dataset. This means that as a learning-based method, scDeepHash can model the latent distribution of a given dataset very well despite the data’s complexity, which helps broaden its application to more general use cases of scRNA-seq data with different characteristics.

#### Inter-dataset annotation accuracy

In realistic scenarios, the source and target datasets are often generated from different experimental platforms by different labs. To evaluate scDeepHash in this realistic setting, we also conducted inter-dataset experiments on four paired source-target human pancreatic datasets: Baron Human, Muraro, Segerstople, and Xin. Simmilar to (Abdelaal et al. [2019]), we tested four combinations by training on three datasets and tested on the remaining one dataset, where the annotation performance can be affected by batch differences among datasets. Aside from the original data, we also tested scDeepHash on the aligned data preprocessed with mutual nearest neighbor method (Haghverdi et al. [2018]) to better group the combined cell data.

Figure 2(b) shows the overall inter-dataset performance of scDeepHash and other competing methods evaluated with the median F1-score. For the original datasets, most state-of-the-art methods perform relatively well on the four combinations of experiments. However, CASTLE exhibits a sharp performance drop when being tested on Baron Human dataset, while scDeepHash performs the best. For the aligned datasets, scDeepHash and SVM perform consistently well across 4 experiments. Other competing methods all encounter a sharp drop or even failure on one or more datasets (like SinglecellNet on Baron Human dataset, scVI on Xin, etc.), which is consistent with the findings reported in (Abdelaal et al. [2019]). Overall, scDeepHash demonstrates robust performance for inter-dataset annotation which makes it a well-suited method in realistic scenarios.

#### Annotation Speed

We cross compare the annotation speed among the state-of-the-art methods and see how they scale when the number of testing samples increases. To keep variables including feature sizes, cell populations, and cell correlation consistent, we choose TM as the candidate dataset and only scale the number of testing samples for variation.

Figure 2(d) shows the big discrepancy regarding the speed capacity of different methods. Other than methods like SingleR, CaSTLe with extremely slow annotation (10k+ seconds for total TM annotation), most methods generally scale well. To annotate the whole TM dataset, scDeepHash only needs less than 3 secs, which is exceptionally fast. This echos the contribution of the fast computation and the fast labeling strategy described in Sec.3.3. As the fastest method, scVI performs annotation a magnitude smaller than that of the second-ranking scDeepHash, but it is at the cost of huge sacrifice on annotation accuracy (Figure 2(d)) since it only uses 1-layer neural network for variational inference (Lopez et al. [2018]). For scDeepHash, most of the annotation time is spent on passing the high-dimensional raw cell data through the 5-layer neural network for high-quality hash code generation.

Overall, when considering accuracy and speed as a whole, scDeepHash performs consistently the best against the rest of the state-of-the-art methods on cell-type annotation.

### 4.2 Performance on Similar-Cell Retrieval

While all the above benchmarking methods are limited to annotation task only (e.g, SVM can only predict cell type in a qualitative fashion and cannot rank the similarity of cells), scDeepHash is able to perform similar-cell retrieval task with exceptional accuracy and speed. Here, we benchmark scDeepHash against another two retrieval-based annotation methods: CellFishing.jl (Sato et al. [2019]) and scmap-cell (Kiselev et al. [2018]). For retrieval-based annotation methods, the annotation accuracy is largely determined by how well the method ranks the similarities between query cells and all the reference cells in the database to retrieve. Thus, retrieval accuracy is essential for retrieval-based annotation methods. To benchmark the retrieval accuracy and speed, we made a trivial modification on their final outputs, from returning a cell label to returning most similar cells in their own database for each query. We use Mean Average Precision (mAP) as the evaluation metric since it is the most standard metric for evaluating retrieval tasks.

#### Retrieval Accuracy

Results in terms of mAP for cell data retrieval are given in Table 2, in which benchmarking is performed on two relatively complex datasets - AMB92 (with abundant cell types) and TM (with large sample size and high feature dimensions). We vary the top *k* retrieved cells from database, where *k* is selected based on the average number of ground truth of all cell population in the cell database, respectively (AMB92 is ~90 on average, and TM is ~600). We observe that scDeepHash achieves the best performance on similar-cell retrieval. Compared with the other two methods, scDeepHash exhibits a sharp mAP increase on both datasets, with more than 20% improvement on AMB92. Meanwhile, as the number of retrieval cells (*k*) increases, both other two competing methods encounter sharp decrease while scDeepHash can maintain a robustly stable retrieval performance. Such robustness to *k* indicates that scDeepHash optimizes the retrieval task and can prioritize the cells in query database from a global perspective, as opposed to the other counterparts that perform much worse in finding the relatively ‘less similar’ ground truth given the query cell. Figure 3 (a) shows retrieval accuracy in Precision curves with respect to different numbers of returned samples (P@N). We can also find scDeepHash outperformed the other compared methods by a large margin.

**Table 2.** **Comparison of retrieval accuracy in mAP** against TM & AMB92 varying on top *k*, where *k* is based on the average number of ground truth of all cell population in TM and AMB92, respectively.

| Methods | AMB92 (mAP) |  |  | TM (mAP) |  |  |
| --- | --- | --- | --- | --- | --- | --- |
|  | @50 | @100 | @200 | @100 | @500 | @1000 |
| scmap-cell | 0.743 | 0.704 | 0.663 | 0.928 | 0.888 | 0.864 |
| CellFishing.jl | 0.742 | 0.697 | 0.649 | 0.946 | 0.915 | 0.897 |
| scDeepHash(Ours) | **0.947** | **0.947** | **0.946** | **0.956** | **0.947** | **0.943** |

**Fig. 3.**
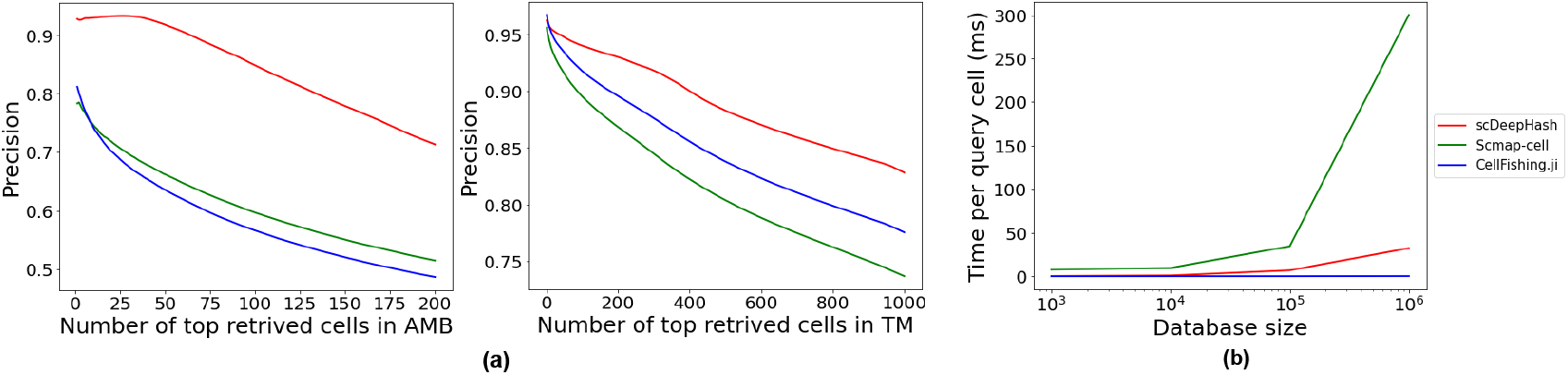
P@N curves and Retrieval Speed Experiment. **(a)**: Retrieval accuracy measured by P@N curves on AMB92 and TM datasets across three methods. Each query cell is used to retrieve the top-*k* most similar cells from their own databases. The average precision across all cell types present in query cells is calculated with respective to each *k*. **(b)**: Retrieval speed measured in millisecond by time per query, with varying database sizes.

#### Retrieval Speed

To guarantee the consistency of potential factors (feature size, cell types, etc), similarly, we use simulated TM datasets (upsampling the original TM to create huge database) for retrieval speed test. The experiment is conducted by retrieving top-10 similar cells given 10k query cells varying on the size of reference database. The average retrieval time per query cell is used as the metric to measure the retrieval speed. As shown in Figure 3(b), scDeepHash clearly outperforms scmapcell, which conducts a linear scan through the entire database for each query cell. Compared with Cellfishing.ji, which uses multi-index hash (MIH) (Norouzi et al. [2013]) to speed up the searching process, scDeepHash performs slightly slower when the size of reference database increases to million-cell level due to the large amount of hash codes that needed to be linearly scanned through in the cell anchor. MIH divides bit vectors into shorter sub-vectors and indexes them separately into ‘hash buckets’. The sub-vectors are later used for k-nearest neighbors search during query time. Even though such design can achieve significant speed improvement, it does sacrifice the retrieval accuracy since searching by sub-vectors is unreliable especially when the size of sub-vector is small.

### 4.3 Ablation Study

#### Different number of hash code bits

We investigate the effects of using different number of bits of the generated hash codes for cell-type annotation. Figure 4 shows the median F1-scores on BaronHuman and TM using hash codes of 32 bits, 64 bits, 128 bits and 256 bits. We observe that increasing number of bits in hash codes doesn’t necessarily lead to performance increase. In fact, as long as we choose the number of bits n such that *M* < 2^*n*^, where *M* denotes the number of cell types, scDeepHash can achieve consistently good performance with varying number of bits. This result further proves that our generated cell anchors well separate distinct cell types and method’s performance is robust to the number of hashing bits.

**Fig. 4.**
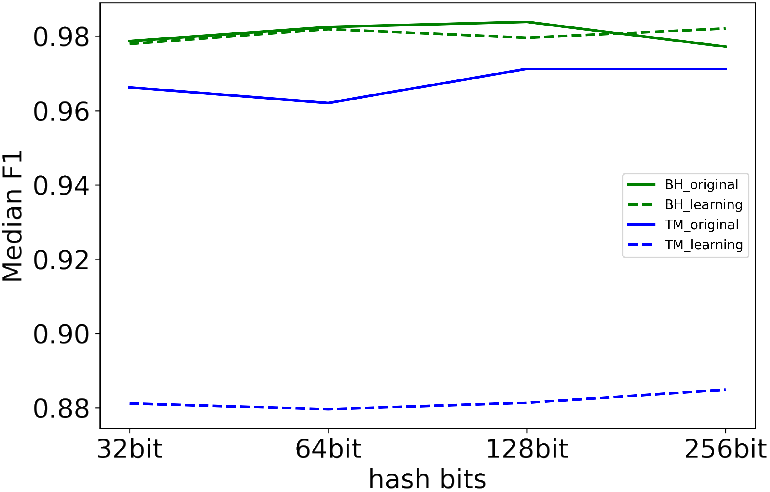
Two ablation studies of scDeepHash on BaronHuman and TM datasets. The first ablation is by varying the number of bits of the generated hash codes. The second ablation is to compare using predefined cell anchors or jointly-learned cell anchors.

#### Learning Cell Anchor

From a biological point of view, not all cell types are equally distant from each other. For example, the ‘similarity distance’ between cell types from the same organ may be smaller than cell types from different organs. Thus, a more intuitive way should be jointly learning the cell anchors rather than predefining cell anchors. To investigate the performance of learning cell anchors, we adopt cell anchor learning in Representative-based Metric Learning (RepMet) (Karlinsky et al. [2019]) to learn cell anchors from features and compare the performance with predefining cell anchors. In RepMet, cell anchor learning is achieved by jointly optimizing a separate fully-connected neural network. This separate neural network takes in vectors **1** as the initialization for all cell anchors and learn the cell anchors that can best separate different cell types iteratively with the hash function training. We apply such a cell anchor-learning strategy and compare the performance of cell-type annotation with the cell anchor-predefining strategy as shown in Figure 4. For relatively simple dataset - BaronHuman, these two strategies achieve comparable performance. However, for more complex dataset TM, we observe a sharp decrease (around 8.3%) on cell anchor-learning one. We hypothesize that the extra binarization to the learned cell anchors results in similarity information loss, so that the learned cell anchors are not distant enough compared with predefined cell anchors, which leads to worse performance in terms of aggregating the corresponding cells of the same type. In conclusion, we choose the predefined cell anchor strategy as our method since it has less computation cost and superior performance.

## 5 Discussion & Conclusion

In this paper, we proposed a novel learning-to-hash methodology for cell-type annotation and similar-cell retrieval. We further quantitatively demonstrated that scDeepHash has a reliable and efficient performance on large-scale datasets compared with other state of the arts. With the ever-increasing scale of scRNA-seq datasets, we believe scDeepHash is a method for the future and it has many directions that worth further exploring.

Our future exploration falls into the following three aspects. First is to incorporate more robust batch correction. In the real-world application, training data and test data are usually generated under different protocols from different labs. Finding the intrinsic latent features and modeling the corresponding distributions is very useful in this case. Second, conveniently incorporating new cell data into the existing scDeepHash model is another interesting topic. With ever-increasing scRNA-seq data, new cell subtypes, or even new cell types, it can be time-consuming and cumbersome to iteratively incorporate novel cell data into the existing model. A worth-discovering topic is to study the incremental hashing methodology (Wu et al. [2019]) or other online-learning methods to see how such capability can be fulfilled in scDeepHash. Third, detection of novel cell types can be very useful for both cell-type annotation and similar-cell retrieval. Scenarios where the true types of query cells are never seen by the underlying model can happen, and thus forcing label assignment or retrieved cells will be unavoidably unreliable. Various methods including scPred (Alquicira-Hernandez et al. [2019]), CHETAH (Kanter et al. [2019]) have already incorporated a similar mechanism into the classification task, while proper confidence boundary is still hard to achieve. We will dive deeper into this topic and see how to enable scDeepHash to reliably detect unknown cell types.

